# Learning the sequence of influenza A genome assembly during viral replication using point process models and fluorescence in situ hybridization

**DOI:** 10.1101/322966

**Authors:** Timothy D. Majarian, Robert F. Murphy, Seema S. Lakdawala

**Affiliations:** Computational Biology Department, University of Pittsburgh School of Medicine; Department of Biological Sciences, University of Pittsburgh School of Medicine; Department of Biomedical Engineering, University of Pittsburgh School of Medicine; Machine Learning Department, Carnegie Mellon University; Department of Microbiology and Molecular Genetics, University of Pittsburgh School of Medicine

## Abstract

Within influenza virus infected cells, viral genomic RNA are selectively packed into progeny virions, which predominantly contain a single copy of 8 viral RNA segments. Intersegmental RNA-RNA interactions are thought to mediate selective packaging of each viral ribonucleoprotein complex (vRNP). Clear evidence of a specific interaction network culminating in the full genomic set has yet to be identified. Using multi-color fluorescence *in situ* hybridization to visualize four vRNP segments within a single cell, we developed image-based models of vRNP-vRNP spatial dependence. These models were used to construct likely sequences of vRNP associations resulting in the full genomic set. Our results support the notion that selective packaging occurs during cytoplasmic transport and identifies the formation of multiple distinct vRNP sub-complexes that likely form as intermediate steps toward full genomic inclusion into a progeny virion. The methods employed demonstrate a statistically driven, model based approach applicable to other interaction and assembly problems.

**Author Summary:** Influenza virus consists of eight viral ribonucleoproteins (vRNPs) that are assembled by infected cells to produce new virions. The process by which all eight vRNPs are assembled is not yet understood. We therefore used images from a previous study in which up to four vRNPs had been visualized in the same cell to construct spatial point process models that measure how well the subcellular distribution of one vRNP can be predicted from one or more other vRNPs. We used the likelihood of these models as an estimate of the extent of association between vRNPs and thereby constructed likely sequences of vRNP assembly that would produce full virions. Our work identifies the formation of multiple distinct vRNP sub-complexes that likely form as intermediate steps toward production of a virion. The results may be of use in designing strategies to interfere with virus assembly. We also anticipate that the approach may be useful for studying other assembly processes, especially for complexes with modest affinities and more components than can be visualized simultaneously.

## Introduction

Influenza A virus, part of the *orthomyxoviridae* family, has a segmented genome of eight distinct viral RNA segments coding at least 11 major proteins and several auxiliary peptides. Most notable of the 11 viral proteins are hemagglutinin (HA) and neurominidase (NA), the canonical *H* and *N* in influenza strain designation. A segmented genome offers potential evolutionary advantages during viral replication, in the form of reassortment. The exchange of genomic material between two distinct viral strains in a co-infected cell often causes a shift within the genome, greatly increasing viral genetic diversity. In only the last century, influenza pandemics of 1957 (H2N2), 1968 (H3N2), and 2009 (H1N1) have emerged through reassortment of at least two viruses [1]. Segmented genomes also come with an inherent mechanistic challenge: ensuring that progeny receive a complete set of genomic segments. Random packaging is possible but comes at the cost of producing an overwhelming majority of progeny that are not viable [2]. Evidence suggests that there exists an active mechanism of selective packaging ensuring progeny viability through efficient and accurate genomic packaging [3, 4].

Within a virion, the viral genome is organized into individual viral ribonucleoprotein (vRNP) complexes, composed of the viral RNA, virally encoded nucleoprotein (NP), and a heterotrimeric polymerase complex made up of PB1, PB2 and PA. The vRNP structure is known classically as a helical panhandle, where the RNA is wrapped around NP with RNA bases exposed and the 3’ and 5’ ends associated with the polymerase complex [3]. We have recently demonstrated that the structure of vRNP is more complex than the classically depicted “beads-on-a-string” schematic. The NP and viral RNA associate in a non-uniform manner, where regions of the RNA are unbound by NP and capable of forming complex structures [5]. Macro-organization of vRNPs within the viral capsid show tight packaging, organized in a ‘7+1’ orientation, a single vRNP center shaft surrounded by seven others (4–6).

Given the need to package one copy of all eight vRNP segments, there is strong evidence for the existence of a selective packaging mechanism prior to budding [3, 6–9]. Intersegmental RNA-RNA interactions have been proposed to mediate selective packaging of all eight influenza vRNP segments. The 5’ and 3’ regions of each segment have been implicated to be essential for efficient packaging and may be critical for the proposed RNA-RNA interactions [3, 10–13]. Previous studies using *in vitro* transcribed viral RNA have successfully demonstrated RNA-RNA binding. These studies also show that a set of eight reproducible interactions can suffice to form a full genome complex [14–16]. However, these studies were performed on viral RNA in the absence of NP which would alter the regions of viral RNA accessible for the proposed RNA-RNA interactions. Recent *in vivo* experiments have also pointed towards RNA-RNA interaction as the driving mechanism for selective packaging [17]. While there is consensus of its existence, a clear network of segment interactions leading to a full genome complex has yet to be elucidated.

We and others have previously used fluorescence *in situ* hybridization (FISH) studies to visualize the intracellular localization of multiple vRNP segments during a productive viral infection [18, 19]. These studies, performed eight hours post infection to capture an initial infection cycle and reduce complication of cytoplasmic vRNP from a subsequent infection, have shown that distinct vRNP segments colocalize within the cytoplasm. Based on our previous studies, we proposed a model whereby vRNP segments are exported from the nucleus, the site of vRNP synthesis, as subcomplexes that undergo further assembly en route to the plasma membrane through dynamic fusion or colocalization events [19]. We also performed a series of binary colocalization comparisons, which would postulate a simple linear vRNP interaction network. However, no such network was identified, suggesting the presence of a more complex vRNP interaction network, including higher order subcomplexes as intermediate steps [19].

The lack of a method to image all eight vRNP segments within a single infected cell has limited our ability to examine the precise spatial relationship between segments. In this study, we developed a model based approach, rooted in point process theory, to quantify vRNP spatial dependence as a metric for RNA-RNA interaction using multi-color FISH images. A spatial point process model is a statistical model that captures the probability that an event will happen at any position within a given geometry. The probability can depend on the spatial distributions of other events or objects, often referred to as covariates or factors. In this context, an event is the observance of fluorescent signal (corresponding to one or more vRNPs) at a given location. We have previously used point process modeling to construct generative models reflecting the spatial dependencies between various punctate cell organelles and other cellular structures, such as the cell and nuclear membranes, microtubules and the endoplasmic reticulum [20, 21]. Related analytical methods have been used to analyze clustering of molecules reflecting spatial dependencies between copies of the same structure [22, 23] Here we use point process models to present a statistically rigorous analysis of intracellular influenza A genome assembly dynamics from a spatial perspective during a productive viral infection.

For an imaged cell, the locations of each vRNP segment can be represented as “realizations” of an underlying probability density specific to that vRNP, calculated over the cytoplasm, and dependent on the spatial proximity to various other structures within the cell. That is, at any given position within the cell, there exists a probability of observing a single segment, of a particular identity, that depends on the positions of cellular organelles and other vRNP segments. By modeling the locational densities of each vRNP, we can therefore learn a dependency network from which likely vRNP interactions are implied.

FISH imaging yields many distinct observations of individual cells, each with multiple vRNP patterns observed. These point patterns, when viewed as realizations of point processes, allow for statistical learning of spatial dependencies over many replicates for each vRNP segment. By defining a set of covariates as the minimum distances between a given segment and other segments observed within the same cell, we produce a set of models that describe the spatial relationship between distinct segments. The extent to which two (or more) vRNPs are dependent upon each other (e.g., likely to be found at nearby positions) is reflected in the likelihood that images of those vRNPs would have been produced by a learned model: high likelihoods signify that the positions of a given vRNP can be predicted well from the others in that model. Model likelihoods can then be seen as a metric of vRNP interaction, with higher likelihoods indicating more probable vRNP association. These likelihoods can then be used to construct the most probable sequence of interactions to form the full genome with methods borrowed from evolutionary tree construction [24].

## Results

### Imaging influenza virus infected cells

To investigate the spatial dependence between vRNP segments, we utilized images from a previous study of infected cells stained for different combinations of vRNP segments using four-color FISH at eight hours post infection [19]. The data set included 14 different probe combinations that covered all pairwise vRNP associations and 32 out of 56 possible triple vRNP combinations (Supplemental Table 1 and 2). Prior to initiating pattern analysis, each image was processed to remove background, isolate individual cells, and find punctate vRNP segments. Figure 1a and b presents representative raw and segmented images with detected points.

**Figure 1:**
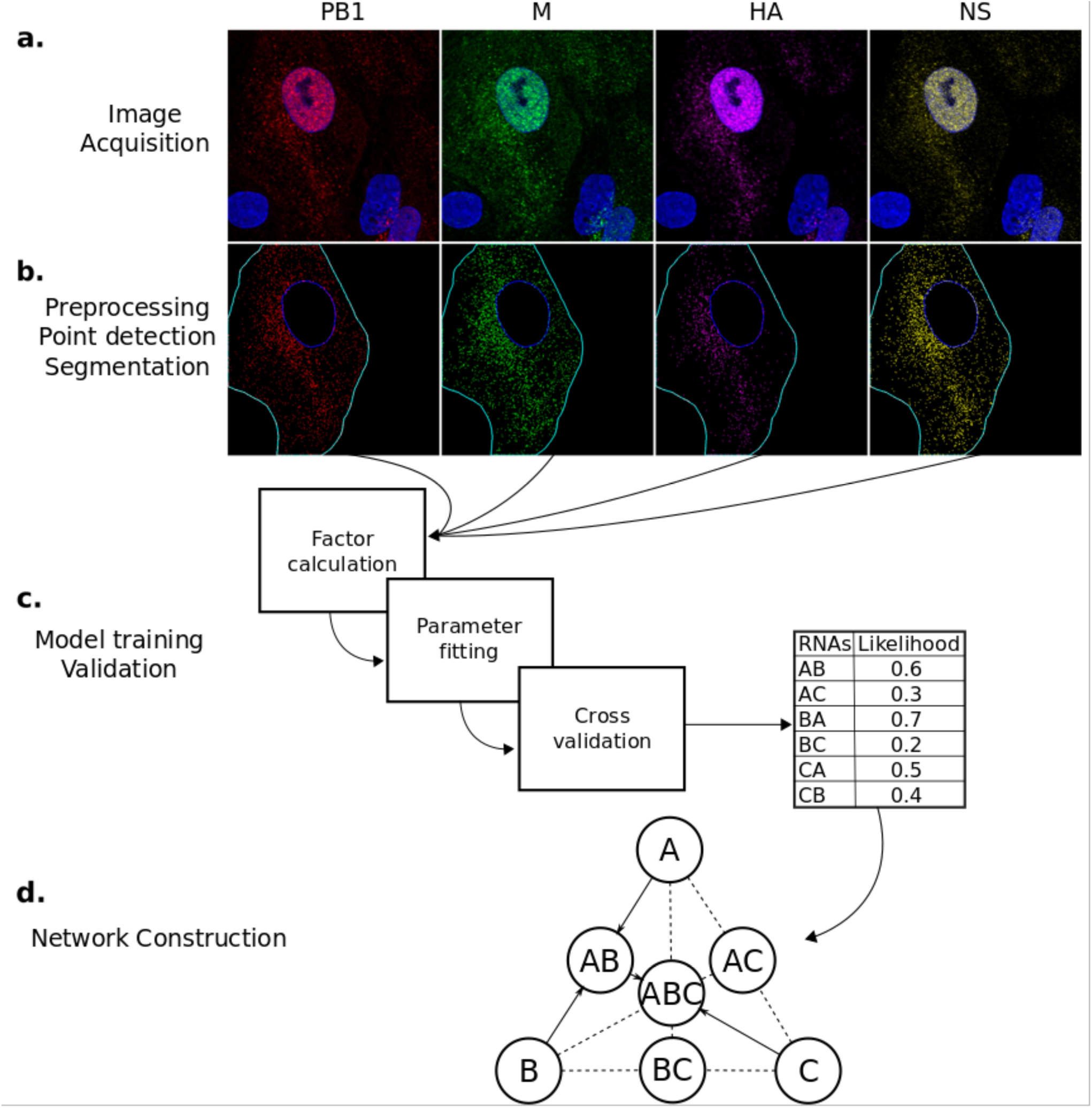
Illustration of the analysis pipeline. (a) Unprocessed fluorescence channels for cells FISH tagged for four vRNA segments and stained with DAPI. (b) Preprocessed image channels after filtering, nuclear and cell membrane segmentation, and point identification. The 3D images in (a) and (b) are displayed as maximum value projections. (c) Factors are calculated for each channel in each image, model parameters are fit, and a likelihood generated. (d) Dynamic programming is used to generate a most likely set of interactions yielding a full genome. Edges are possible interactions, solid lines indicate edges that are part of the most likely set of interactions. In this toy example, the most likely set of interactions is A and B binding to form AB, and then C binding to AB.

### Assessing randomness of the spatial distributions of vRNP

We began by determining the extent to which the individual vRNPs could be considered randomly distributed throughout the cell, since our search for spatial dependencies between different vRNPs would be illogical if their positions were all random. We defined the null hypothesis as complete spatial randomness, as would be exhibited by a homogenous Poisson process, and estimated a p-value for this hypothesis using Monte Carlo methods. Briefly, for a given vRNP in a particular cell, we started by segmenting the cytoplasm and counting the number of points observed within. We then simulated (generated) many random point configurations, of the same number of points, within the segmented cytoplasm. We calculated a test statistic for each random configuration (see Methods) and used these to assign a p-value to the test statistic for the observed pattern for that cell; these were averaged for each vRNP. As shown in Table 1, every vRNP segment showed significant deviation (at the p < 0.05 level), with low variance, from a spatially random distribution. This indicates the presence of a spatial trend and some dependency on cellular structures for their location.

**Table 1:** Monte Carlo p-values and variances for the hypothesis that the spatial distribution of each vRNA segment is distributed by a homogeneous Poisson process.

|  | Mean p-value | Variance |
| --- | --- | --- |
| PB2 | 0.0034 | 0.00011 |
| PB1 | 0.00524 | 0.00034 |
| PA | 0.00467 | 0.0002 |
| HA | 0.00671 | 0.00015 |
| NP | 0.01185 | 0.00036 |
| NA | 0.00249 | 0.00003 |
| M | 0.00005 | 0 |
| NS | 0.00306 | 0.00007 |

### Modeling cell and nuclear membrane dependence

Given that the localizations of vRNPs are not random, we sought to determine if some relationship to the cell and nuclear membranes could explain the spatial trend of vRNP segments. Three covariates were defined for each position in the cell (minimum distance to the cell membrane, minimum distance to the nuclear membrane, and the minimum ratio of cell to nuclear distance for each point) and models were created for each vRNP using each of the covariates separately. We estimated the accuracy of each model by cross validated likelihood calculation (see Methods), where values closer to 0 are more likely. It is important to note that, because actual likelihoods were calculated, they can be compared between models of different complexity.

Dependency on the cell membrane produced the lowest likelihood (worst) models relative to all others (Table 2) for all vRNP segments. Congregation of vRNP to the apical cell membrane is a potential skewing factor, since there will be a high concentration of all segments at this cellular location. The many observed points within close proximity to the membrane potentially caused low fitted probability of observing other points closer to the nucleus. Cell shape differences combined with lower resolution in the z-dimension may have also influenced these models.

**Table 2:** Cross validated negative log-likelihoods for models dependent on distance to different reference locations.

|  | Cell Model | Nuclear Model | Cell+Nuclear Model |
| --- | --- | --- | --- |
| PB2 | 2.5547 | 0.4153 | 0.9158 |
| PB1 | 2.3908 | 0.408 | 1.2574 |
| PA | 2.4228 | 0.4643 | 1.2352 |
| HA | 2.443 | 2.3928 | 4.396 |
| NP | 2.4885 | 2.4584 | 4.5447 |
| NA | 2.5019 | 2.4372 | 2.1754 |
| M | 2.4871 | 2.1294 | 3.0799 |
| NS | 2.3906 | 0.6666 | 1.4624 |

For most vRNPs, models incorporating nuclear distance showed higher likelihoods than those using cell membrane distance (Table 2). Some vRNP segments, such as PB2, PB1, PA and NS, showed greater dependency on nuclear distance than the other segments. The ratio of cell membrane to nuclear distance generally produced models better than cell membrane alone, but they were worse than nuclear alone in all cases except NA (Table 2). Thus nuclear distance provides the most cogent model for dependence of vRNP localization on a cellular structure.

### Inter-segment dependence in pairs, triplets, and quadruplets

Every pair of vRNP segments was observed in at least three cells, allowing for modeling of vRNP-vRNP dependence among all segment pairs. For each pair of vRNP segments, two models were trained, switching the dependent (primary) and independent (secondary) patterns. Since the minimum inter-pattern distance measure is not commutative, the two models need not be the same. More simply, a given vRNP location may depend on its proximity to another vRNP, but the converse is not necessarily true. This can be observed when one vRNP segment is found in two distinct locations, and a second vRNP segment is found in only one of those locations. The second vRNP segment would be observed to have a high likelihood for being predicted from the first, but the first would not have a high likelihood of being predicted from the second.

In all cases, pairwise inter-pattern distance models (Table 3) for a given vRNP had higher likelihoods than those produced for that vRNP with only the cell and/or nuclear membrane factors (Table 2). Some pairs showed higher likelihoods than others, such as PB1-NP, PB1-M, PA-NP, HA-PB1, and NP-PB1. These pairings also showed relatively low variance in cross-validated likelihood estimates (Supplemental Table 3). Many of the highest likelihood vRNP pairs are observed between segments encoding the viral polymerase components (PB2, PB1, PA and NP). Given the dependence between these proteins both structurally and functionally [25–27], it is possible that these segments evolved a dependence upon each other to ensure their joint packaging in a virion (a possibility that would require extensive additional work to examine). Figure 2 highlights the pairwise models in Table 4 by illustrating vRNP pairs that are below 0.23 log likelihood. Overall, vRNPs showed the greatest dependence on PA and PB1, with the most connections between these two segments and the others. The least dependence was observed for NA and NS vRNP segments.

**Table 3:** Cross validated negative log-likelihoods for pairwise interaction models. The row label indicates the independent (secondary) variable, the vRNP upon which a model predicting the column label (the dependent or primary variable) was constructed.

|  | PB2 | PB1 | PA | HA | NP | NA | M | NS |
| --- | --- | --- | --- | --- | --- | --- | --- | --- |
| PB2 |  | 0.3705 | 0.2694 | 0.282 | 0.3146 | 0.5605 | 0.2146 | 0.3926 |
| PB1 | 0.2398 |  | 0.2018 | 0.2044 | 0.1186 | 0.4015 | 0.1701 | 0.2905 |
| PA | 0.2276 | 0.2217 |  | 0.2284 | 0.1502 | 0.3222 | 0.2816 | 0.2315 |
| HA | 0.3334 | 0.1266 | 0.1838 |  | 0.2787 | 0.232 | 0.2623 | 0.2845 |
| NP | 0.3438 | 0.1805 | 0.1942 | 0.2895 |  | 0.2529 | 0.275 | 0.3411 |
| NA | 0.3191 | 0.324 | 0.3093 | 0.2947 | 0.2742 |  | 0.2747 | 0.2982 |
| M | 0.3287 | 0.2236 | 0.2832 | 0.2803 | 0.2975 | 0.2897 |  | 0.2848 |
| NS | 0.3427 | 0.2888 | 0.2292 | 0.2838 | 0.325 | 0.3436 | 0.2841 |  |

**Figure 2:**
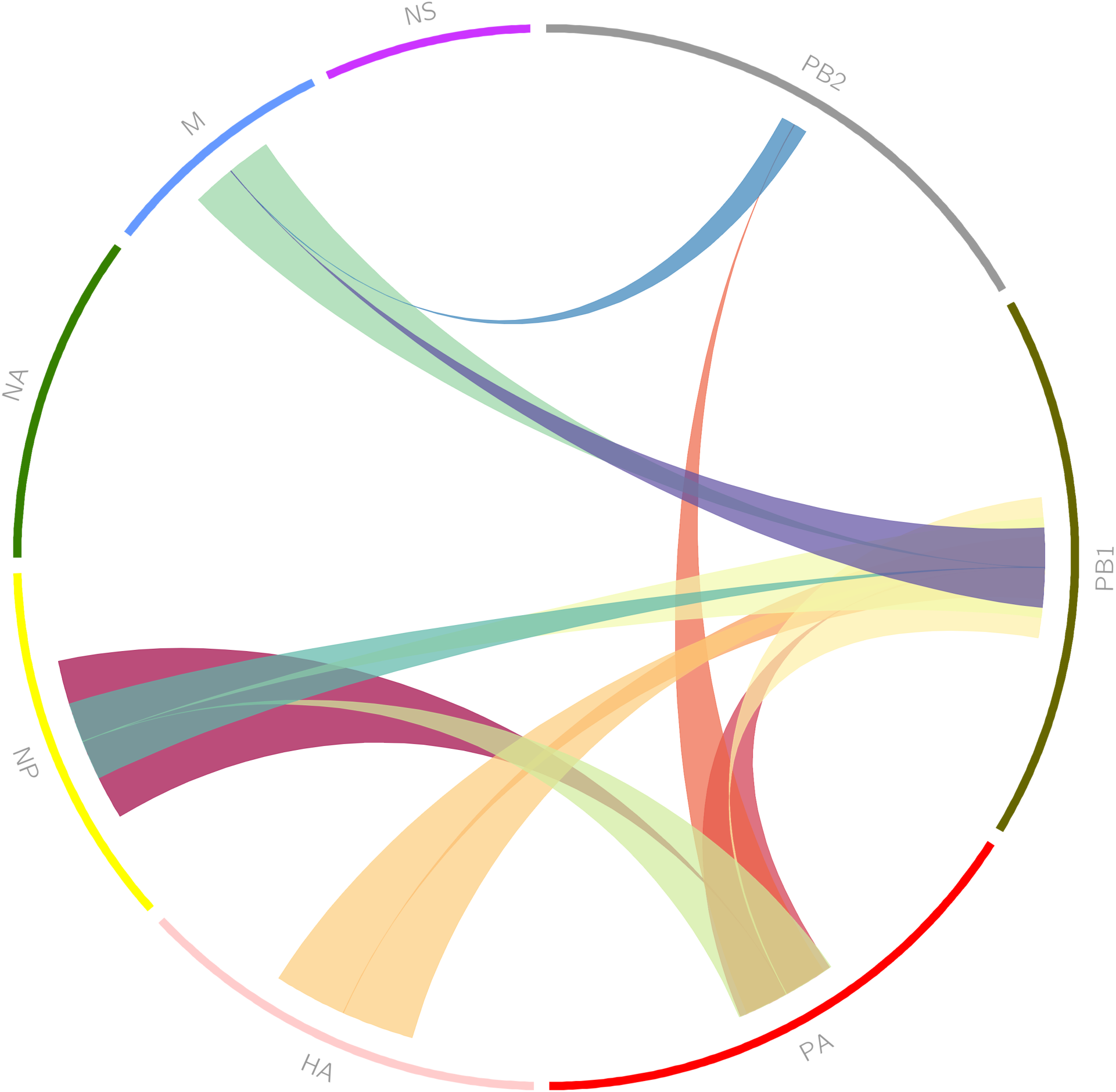
Pairwise interactions learned from MDCK cell images. Each vRNA segment is arranged around the perimeter. Bands connect vRNA segments that interact with high likelihood. The position of the base of each band indicates the independent (secondary) variable and the larger the width of the base the more likely the interaction. Only interactions whose log likelihoods are below 0.23 from Table 4 are shown.

**Table 4:**
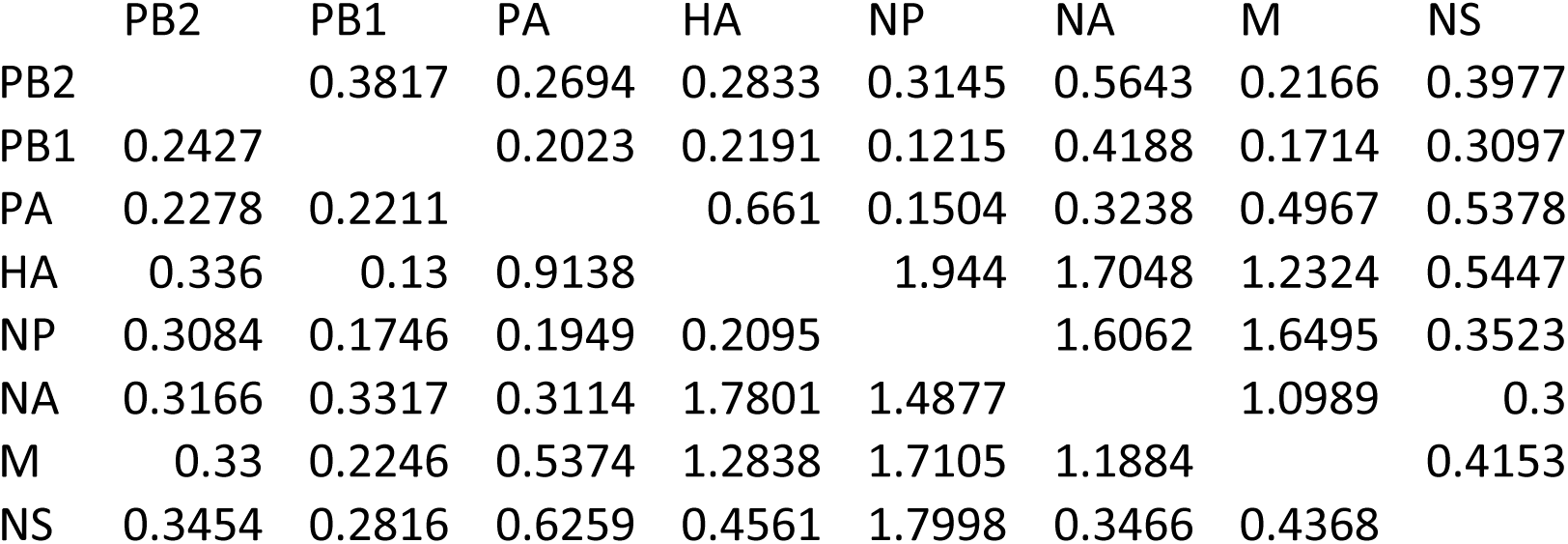
Cross validated negative log-likelihoods for pairwise interaction models that also include nuclear distance. The row label indicates the independent (secondary) variable, the vRNP upon which a model predicting the column label (the dependent or primary variable) was constructed.

Extending the pairwise models, nuclear distance was added as another covariate. Minor improvements in model likelihood were seen in some models (Table 4) with others showing decreased likelihood. Overall there was an increase in the variance of our estimates (Supplemental Table 4). Overfitting to nuclear distance may be the cause of this increased variance.

Thirty-two of 56 possible triplet vRNP segment complexes (referred to as triplets) were present within the dataset analyzed and could therefore be modeled within the same cell. This process allows for modeling of complex vRNP relations containing more than two vRNP segments. Three models were trained for each observed triplet, one for each vRNP depending on the other two. The highest likelihood model was chosen. Model likelihoods tended to be higher than that of either pair model for a given vRNP (Table 5). We observed that the most likely triplet models for each vRNP included four of the five most likely vRNP pairings observed in pair likelihood models (Table 3), (namely PB1-NP, PA-NP, HA-PB1, and NP-PB1). However, not all of the most likely pairwise associations (e.g., PB1-M) were observed in the most likely triplet models, highlighting that models based solely on pairwise comparisons might not accurately represent larger order vRNP complexes. Model likelihoods for all triplets observed can be found in Supplemental Table 5.

**Table 5:**
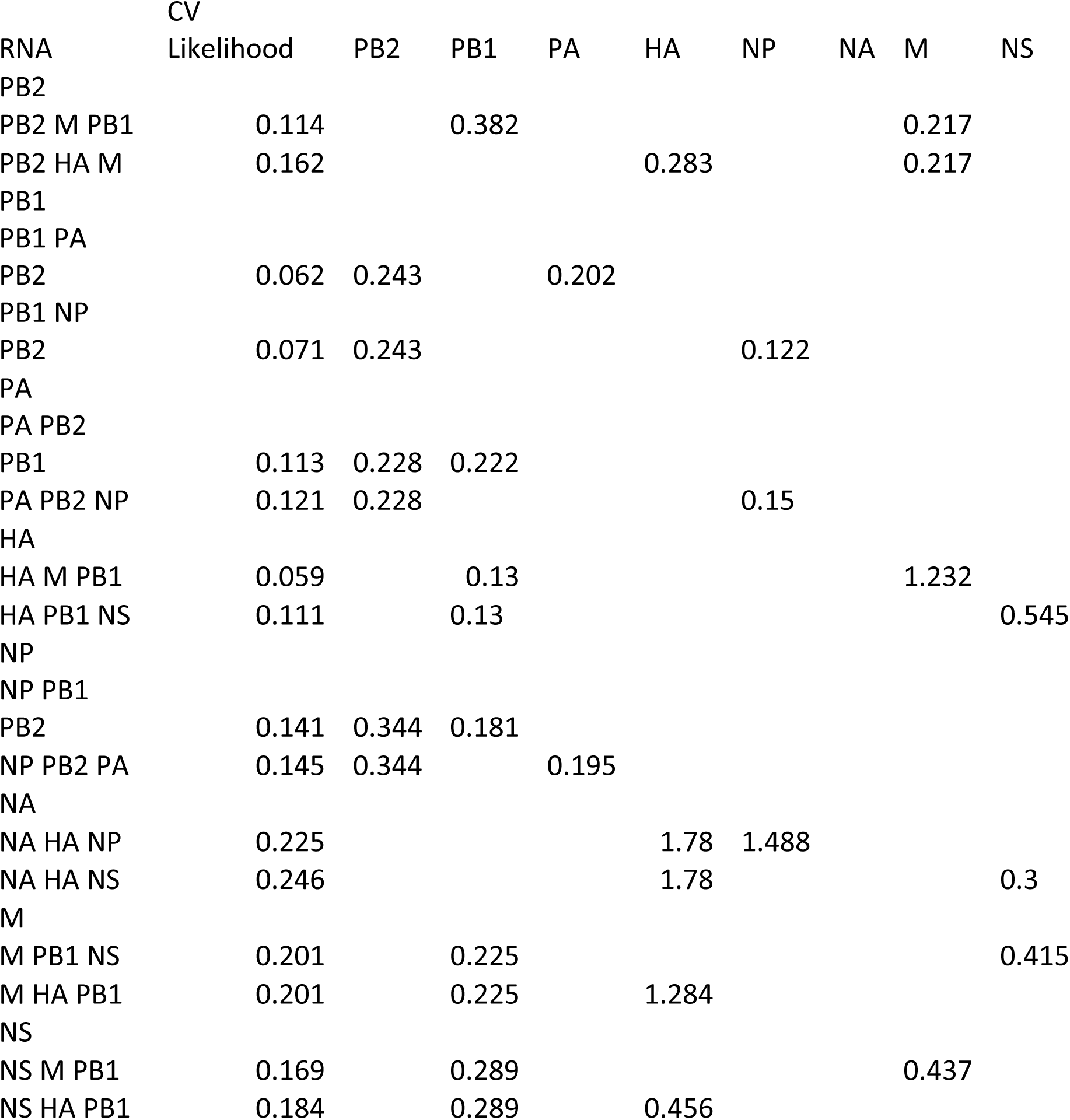
Cross validated negative log-likelihoods for the top two triple models with each vRNA segment as the primary pattern. Pairwise model negative log-likelihoods for the members of each model are shown in the right columns.

A small subset of all possible four vRNP segment complexes (referred to as quadruplets) were contained within the dataset and models for predicting one of the four from the other three were also constructed. Again, considerable overlap is seen between the most likely quadruplet models and both triplet and pair models (Table 6), such as PB2-PB1-PA and PB1-PA. The consistency in vRNP composition between observed likely quadruplet, triples, and pairs further validates the spatial dependence of vRNP segments and the presence of vRNP subcomplexes as influenza A assembly intermediates. Model likelihoods for all quadruplets observed can be found in Supplemental Table 6.

**Table 6:**
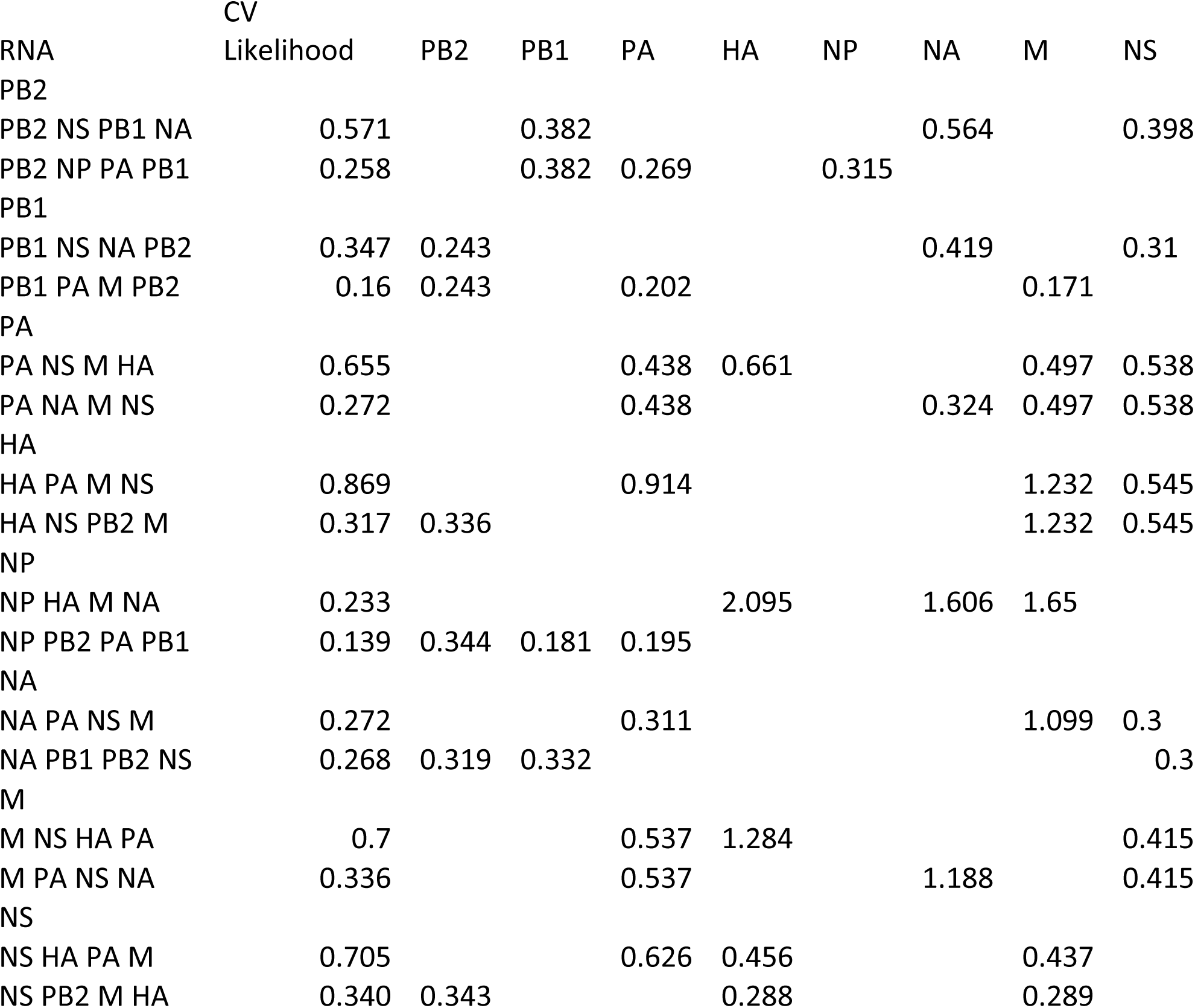
Cross validated negative log-likelihoods for the top two quadruple models with each vRNA segment as the primary pattern. Pairwise model negative log-likelihoods for the members of each model are shown in the right columns.

| RNA | CV<br>Likelihood | PB2 | PB1 | PA | HA | NP | NA | M | NS |
| --- | --- | --- | --- | --- | --- | --- | --- | --- | --- |
| PB2 |  |  |  |  |  |  |  |  |  |
| PB2 NS PB1 NA | 0.571 |  | 0.382 |  |  |  | 0.564 |  | 0.398 |
| PB2 NP PA PB1 | 0.258 |  | 0.382 | 0.269 |  | 0.315 |  |  |  |
| PB1 |  |  |  |  |  |  |  |  |  |
| PB1 NS NA PB2 | 0.347 | 0.243 |  |  |  |  | 0.419 |  | 0.31 |
| PB1 PA M PB2 | 0.16 | 0.243 |  | 0.202 |  |  |  | 0.171 |  |
| PA |  |  |  |  |  |  |  |  |  |
| PA NS M HA | 0.655 |  |  | 0.438 | 0.661 |  |  | 0.497 | 0.538 |
| PA NA M NS | 0.272 |  |  | 0.438 |  |  | 0.324 | 0.497 | 0.538 |
| HA |  |  |  |  |  |  |  |  |  |
| HA PA M NS | 0.869 |  |  | 0.914 |  |  |  | 1.232 | 0.545 |
| HA NS PB2 M | 0.317 | 0.336 |  |  |  |  |  | 1.232 | 0.545 |
| NP |  |  |  |  |  |  |  |  |  |
| NP HA M NA | 0.233 |  |  |  | 2.095 |  | 1.606 | 1.65 |  |
| NP PB2 PA PB1 | 0.139 | 0.344 | 0.181 | 0.195 |  |  |  |  |  |
| NA |  |  |  |  |  |  |  |  |  |
| NA PA NS M | 0.272 |  |  | 0.311 |  |  |  | 1.099 | 0.3 |
| NA PB1 PB2 NS | 0.268 | 0.319 | 0.332 |  |  |  |  |  | 0.3 |
| M |  |  |  |  |  |  |  |  |  |
| M NS HA PA | 0.7 |  |  | 0.537 | 1.284 |  |  |  | 0.415 |
| M PA NS NA | 0.336 |  |  | 0.537 |  |  | 1.188 |  | 0.415 |
| NS |  |  |  |  |  |  |  |  |  |
| NS HA PA M | 0.705 |  |  | 0.626 | 0.456 |  |  | 0.437 |  |
| NS PB2 M HA | 0.340 | 0.343 |  |  | 0.288 |  |  | 0.289 |  |

One of our major goals was to attempt to extrapolate higher order relationships from the pair, triple and quadruplet models. To project how accurately this might be done, we compared our observed triple and quadruple models with predicted models calculated from any of the four pair models (vRNP pairs (Table 3), vRNP pairs + nucleus (Table 4), vRNP pairs +cellular membrane, and vRNP pairs +nuclear/cell membrane ratio) (Figure 3a). We also compared observed quadruplet likelihoods to those predicted using both the pair and triple models, similar to the pair models any of the 4 models including the various cellular features were included (Figure 3b). Most of our observed triples and quadruplets could be predicted with modest accuracy from the pair models but the overall r^2^ value was only 0.302. This was due to inaccuracy in predicting the triple and quadruple clusters containing the PB2-NA pair with either NS, PB1 or both. These combinations were predicted to occur much more often than was observed in the experimental data sets. The pairwise likelihood of NA with PB2 is >-0.6, which is consistent with the observed likelihood and may be the driving force behind the score compared to the other pairwise interactions. Analysis of the predicted triples and quadruples after removing the PB2-NA containing clusters resulted in a r^2^ value increases to 0.642 (we cannot justify this removal on a statistical basis but rather on the observation that the combinations with the poorest predictions all involved the same pair). Predicting quadruples from pairs and triples resulted in a r^2^ value of 0.450, and with removal of the same quadruple it rose to 0.714. Overall, these data support the notion that pairwise point process models can be used to predict higher order complexes with moderate accuracy, and that prediction accuracy increases if higher order models are used.

**Figure 3:**
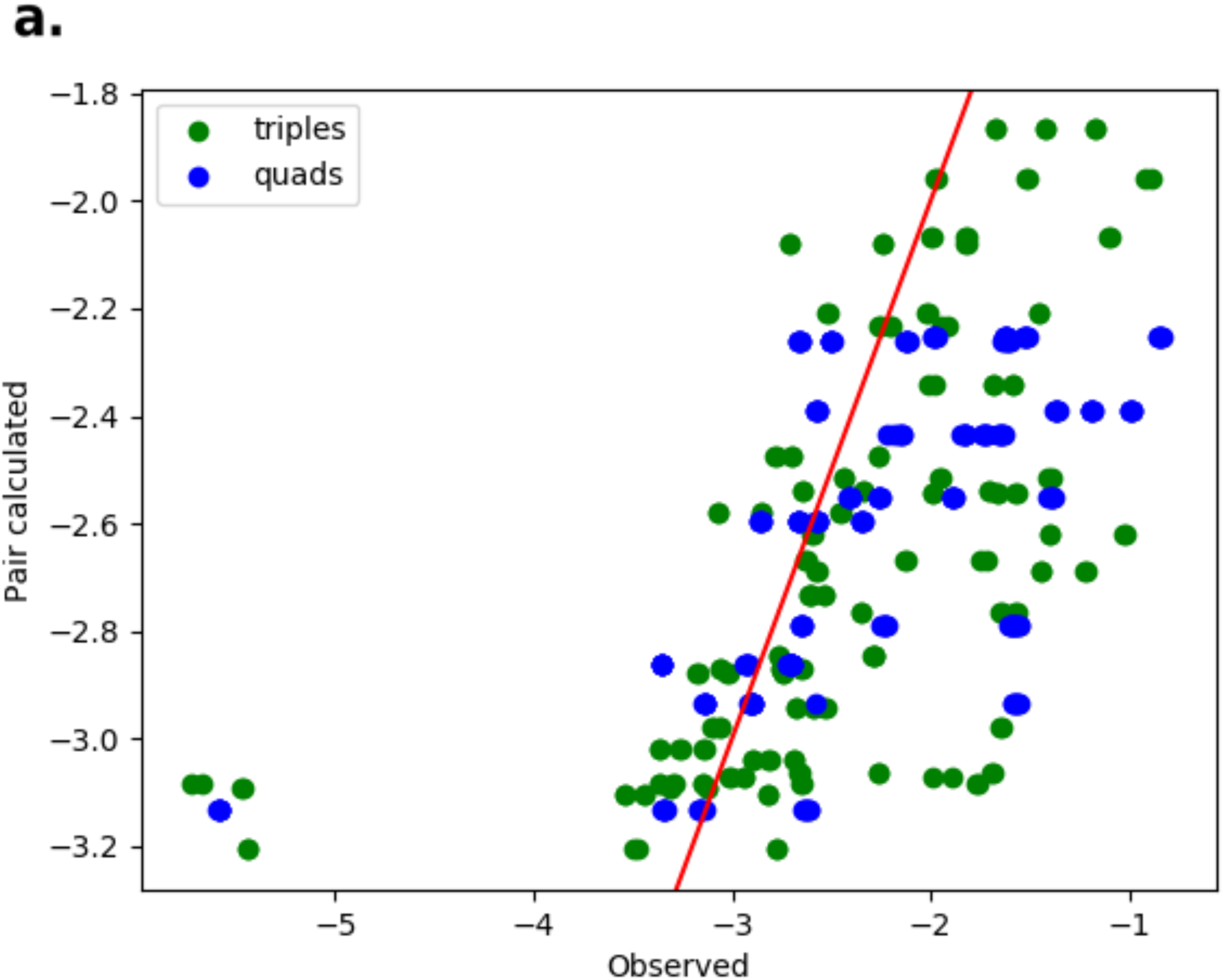
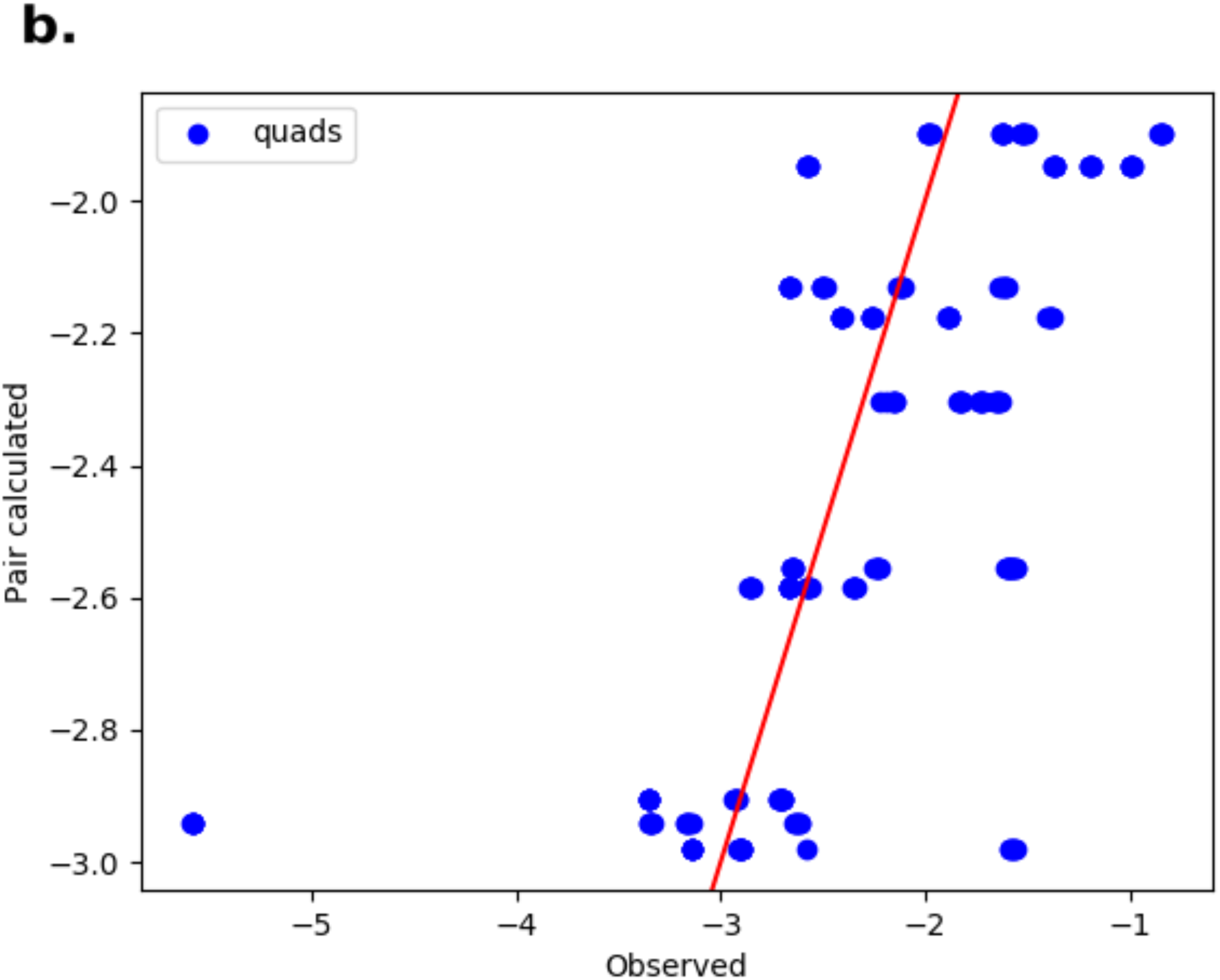
Comparison of composite and observed model likelihoods for triplet and quadruplet models. To ensure that the recursion in our dynamic programming scheme accurately predicts model likelihoods, we compared triplet and quadruplet observed likelihoods to those generated from only pairwise models (that include all four possible models: vRNP pairs alone, vRNP pairs + nucleus, vRNP pairs + cell membrane, vRNP pairs + nuclear/cell membrane ratio) (a). We also compared observed quadruplet likelihoods to those generated from pairwise and triplet models (using all 4 possible models) (b). Ideally, all points would fall on the red line, matching observed and predicted values perfectly. The cluster in the lower left corner represent interactions that are predicted to occur with high likelihood but in reality, do not. All points within this cluster contain the same four vRNA segments (PB1, PB2, NA, NS). Exclusion of this cluster from our analysis increases the association between predicted and observed triplets and quadruplets.

### Interaction network construction

Given these results, we sought to create a model of the set of vRNP interactions (complexes) that occur at all steps in the formation of a complete viral genome. To do this, we represented all possible interactions between vRNP segments as a weighted, directed, acyclic graph with nodes labeled by each unique, unordered set of vRNP segments. A path from all single nodes to the root represents a set of vRNP interactions resulting in a full genome complex (see Methods). For illustration, if all eight vRNP segments directly formed a complete set without forming any intermediate complexes (a highly unlikely scenario), this would be represented by weights of one for the connection between each vRNP and the complete vRNP supramolecular complex containing all eight segments and weights of zero for all other complexes. Such a scenario would result in a tree with the root connected to each of the leaves

In our analysis, we estimated the weight for each possible intermediate vRNP complex assembly subcomplex to contain at least two and up to eight segments based upon either the observed or predicted likelihoods (see Methods). Dynamic programming, an efficient method for recursively finding the highest scoring path to a given node, was then used to find the most probable path within the graph using the highest likelihood trained model for a given nodes from those with or without cell and/or nuclear features (Figure 4a). The resultant tree produces two distinct clusters HA, M, NA, NS (hereafter referred to as C1) and PB1, PB2, NP (C2) that merge with PA as a final step towards forming the full genome. This network suggests three steps to form a complete set of all eight vRNP segments: 1) formation of C1, 2) formation of C2 and 3) addition of PA to C1 and C2 to form the final product.

**Figure 4:**
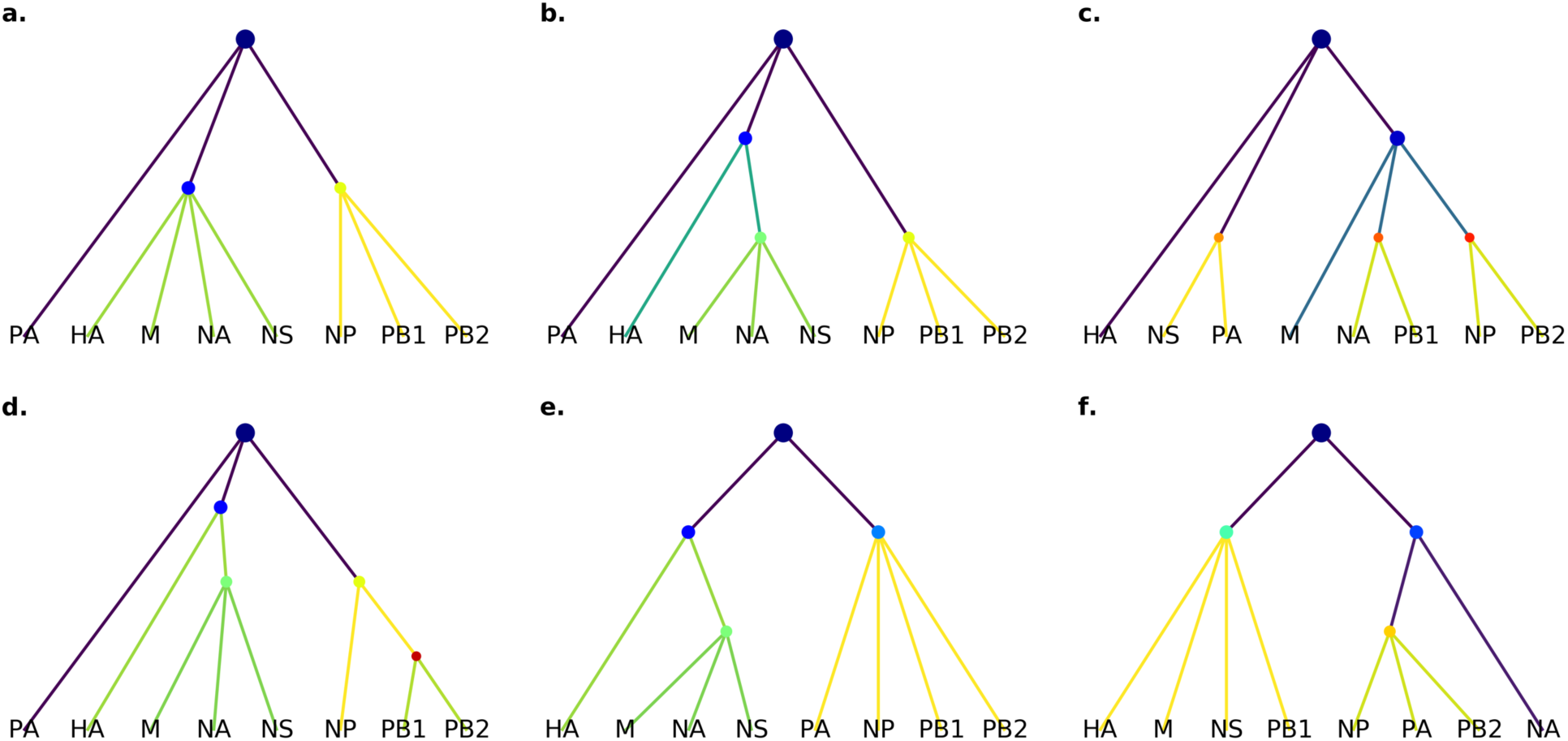
Constructed interaction networks. Using dynamic programming, we constructed the single most likely set of interactions yielding a full genome. First, we used all model likelihoods with and without nuclear and cell features (a) to build a master tree. Hotter edge colors correspond to more likely interactions. Since the images contain only a subset of the possible triplets and quadruplet, we assessed how much this affected the master tree by removing all quadruplets (b) or both quadruplets and triplets (c). To test the stability of the master tree, trees were constructed with random noise added to each observed model likelihood before performing the dynamic programing. Multiple noise levels were tested up to 1/2 of the mean model likelihood with the noise level being the standard deviation of a Gaussian centered at 0. For each level of noise, network construction was performed 100 times. Noise up to 1/4 mean likelihood always resulted in (a). Increasing noise levels yielded (a) on 40% to 60% of constructions. Given the two observed clusters in the master tree, we next asked whether the composite likelihood function could recapture their interactions without actually observing them. Removing all models involving HA, M, NS, NA (C1) and NP, PB1, PB2 (C2) yielded (d). Adding noise up to 1/4 mean likelihood and removing both clusters also generated (d). Removal with higher levels of noise generated (e) for 27% of constructions. In all stability analyses with noise greater than 1/4 mean likelihood, (f) was generated with frequency between 5% and 15%.

To assess whether a particular vRNP localization dataset was driving this network, we performed a series of network analyses with data excluded. First, the contribution of observed quadruplets and triplets was assessed by excluding observed quadruplets (Figure 4b) and without observed quadruplets and triplets (Figure 4c). Interestingly, exclusion of all observed quadruplets did not drastically alter the network, with both C1 and C2 still being present, although formation of C1 required two steps. However, networks built only with observed pairs (excluding triplets and quadruplets) resulted in a quite different network lacking larger subcomplexes (Figure 4c), reflecting the fact that pairs were only able to make approximate estimates of higher order interactions

Given that the dynamic programming approach yields only a single most likely tree, we sought to assess the robustness of the inferred tree to potential inaccuracy in the estimated dependencies. We therefore assessed the stability of the tree by adding various amounts of random noise to the estimated likelihoods (see Methods). Surprisingly, the addition of noise of up to a quarter of the mean log-likelihood had no effect on the network found, yielding the same network (Figure 4a). As noise levels increased to half of the mean log-likelihood, one other tree was generated (Figure 4f). This demonstrates the stability limit of the construction but notably this revised tree still contains clusters that resemble the original. If the original tree was based on biased likelihoods from imaging artifacts, the addition of noise would have resulted in an altered tree network at lower noise levels and with higher frequency, but since this did not occur, we are confident in the accuracy of our original likelihood estimations.

Finally, to exclude the possibility that datasets directly measuring C1 and C2 (reactions L and B, respectively) were biasing the network towards inclusion of those sets of vRNPs, we performed a network analysis of a dataset where the measured likelihoods of these two clusters was excluded. Removal of data derived from images of the sets of vRNPs in C1 and C2 (Figure 4e) yielded networks that resemble Figure 4a, indicating that even in the absence of data from these clusters, similar clusters will form during assembly. Removal of each cluster alone was recapitulated by lower order interactions. Removal of C1 [HA, M, NA, NS] data resulted in the same tree presented in figure 4b, where HA, M, NA, NS form in two steps from a triplet and a single vRNP segment. Exclusion of C2 [NP, PB1, PB2} result in a tree similar to figure 4d, where C2 cluster forms from a pair and a single segment (Figure 4d). Removal of data was also combined with the addition of noise to further test stability. Predominantly, the same networks were generated with and without noise. As noise levels increased in the exclusion constructions, a tree with HA, M, NS, *PB1* in place of C1 and NP, *PA*, PB2 in place of C2, where PB1 and PA were not observed in the C1 or C2 original clusters respectively (Figure 4f). Together, these constructed networks suggest a primary interaction scheme that can be further experimentally tested and the importance of triplet and quadruplet observations in predicting higher order vRNP complexes. Other, very low frequency trees that were generated in the presence of high levels of noise are located in the reproducible research archive.

## Discussion

Influenza A vRNP segments selectively assemble within an infected cell to produce progeny virions containing one copy of all eight segments. The mechanism driving selective assembly is still largely unknown, but RNA-RNA interaction between the segments has been proposed [14–16]. In this study, we have modeled the *in vivo* spatial dependencies of influenza A vRNP segments using multi-color fluorescent *in situ* images to generate possible vRNP-vRNP interaction networks and propose a new perspective on genome packaging during viral replication. By utilizing a rigorous statistical framework, we have extended the efforts of previous groups and present a novel method for construction of vRNP interaction networks based on their precise spatial information within actively infected cells. Using spatial proximity as a proxy for physical interaction, point process modeling in this study suggests clear spatial dependencies between certain vRNP segments, and confirms our previous observations of subcomplex formation during cytoplasmic transport (18).

Previous studies using *in vitro* transcribed RNA suggest multiple interactions between vRNP segments are expected [3, 6, 14, 16, 28]. Similarly, the modeled likelihoods show potentially multiple interactions for each vRNP segment. Electron tomography studies have revealed a conserved ‘7+1’ supramolecular structure within the viral interior, where a center shaft is surrounded by seven remaining segments [3, 6, 28], demonstrating an ordered process in genome assembly.

Our results suggest two candidates for the center shaft or ‘master segment’: PA and PB1. From the six constructed networks, PA is the most variable in that it is least often paired early in assembly but, with the addition of noise and exclusion of data, may interact with both primary observed clusters. This would be expected if a segment was evenly dependent on most others, a possible signature of a core segment. PB1 also behaves similarly but to a lesser degree. Both PB1 and PA show the highest average pairwise model likelihood over all other segments.

A few vRNP pairs were seen often in the most likely pairwise, triplet, and quadruplet models and in the final constructed network. PB1-PB2 and HA-M, show commutative relationships, appearing in one of the top most likely triplet and quadruplet models for each vRNP within the pair. Some non-commutative dependencies were also seen, most notably in NP depending on PB2 in both triplet and quadruplet models with little PB2-NP dependence.

By using dynamic programming, we were able to generate the most likely interaction network based on all models. While this network was surprising robust to the addition of noise to the model likelihoods, subtle variations in the vRNP interaction network were observed with the exclusion of higher order datasets, suggesting that there may exist multiple interaction pathways all acting at once in a single cell. Some of the variation in our results may also be due to the presence of some lower specificity vRNA-vRNA interactions also occurring during genome assembly. This would explain some of the highly likely pairwise interactions observed that were not included in the final interaction network, such as PB1 and HA. In addition, vRNP networks could also change over the course of a viral infection when packaging fidelity may be compromised [29]. Our current data set only considers one time point, eight hours post infection, which represents the time of initial virion release from infected cells, and would miss alternate networks present at later time points. Therefore, the analysis presented here may only capture a snapshot of a dynamic assembly process that could change throughout an infection. A system with multiple possible interaction networks could be potentially advantageous in both ensuring packaging of a full genome in viral particles and in future reassortment events. For example, reassortment of one vRNA segment during a coinfection may decrease the binding efficacy necessary for a single interaction network but a second network may serve to rescue viral viability while also increasing genetic diversity.

With this, we also point out the consistent presence of the PB2, PB1 and NP cluster. Since these three proteins function together, with PA, to form the viral polymerase, which promote vRNA replication and transcription, this cluster is potentially important. Viruses capable of packaging complementary polymerase segments into progeny virions will have a fitness advantage compared to viruses with polymerase mismatches. This phenotype has been observed in experimental reassortment experiments between 2009 H1N1 pandemic and seasonal H3N2 viruses[25, 27].

More generally, the computational pipeline presented may function as a useful tool in elucidating spatial dependency over a wide range of biological phenomena observable through microscopy. As we have demonstrated, point process models can capture key aspects of point distributions while retaining a basis in probability. Network construction synthesizes the results derived through modeling into a cogent, most-likely set of spatial interactions or dependencies.

Future work in spatial modeling of influenza A packaging should incorporate a temporal dimension through live cell imaging. Point process models have been adapted for spatiotemporal modeling [30]. Incorporation of intracellular markers important for influenza A vRNP transport may increase the accuracy of models. Influenza A vRNP transport from the nucleus, the site of vRNP synthesis, to the plasma membrane is a complex process utilizing a variety of host proteins [31]. Rab11A-containing vesicles are thought to be the primary mode of transport although there is evidence that a Rab11A-independent mechanism exists [32–34]. A recent study has implicated the ER to mediating transport of vRNP segments through anchored Rab11A proteins [35]. Novel methods in fluorescent multiplexing [36] present an exciting opportunity for observing many cellular structures which will provide a holistic image the spatial dependence of vRNP segments upon subcellular structures and to each other.

## Methods

### Multi-color fluorescent in situ hybridization images

This study used previously published multi-color FISH images from [19] that were generated at the National Institutes of Health. Briefly, MDCK cells were infected with recombinant WSN/1933 H1N1 for 8 hours and then fixed and stained with FISH probes, obtained from Biosearch technologies, against four distinct vRNP segments. DAPI was included as well to label DNA. Multi-color FISH samples were imaged on an Leica SP5 white light laser to ensure spectral separation of the five colors (Dapi, Alexa 488, Quasar 570, Cal Fluor Red 590, Quasar 670). The specificity of the probes and spectral separation between fluorophores was previously confirmed [19].

### Image preprocessing, cell segmentation and point detection

All image preprocessing was performed in MATLAB, v. 2015a. Prior to point detection and segmentation, image noise was removed by convolution with a Gaussian mask. Each image channel was denoised individually. As described in [19], each image was captured with 0.17 um z-step size spanning the entire cell volume, defined for each cell using both the nuclear and FISH staining to define the apical cell membrane. Each image was captured with a pixel size of ∼50×50×168 nm.

Individual vRNP segments were identified by finding connected areas of signal within the denoised image. The center of each object was taken as the maximum intensity pixel within, yielding a set of point coordinates. Note that due to the thickness of the z sections, the apparent distance between points may be an underestimate of the true distance, but this effect is expected to average out when considering many points.

Since there was no fluorescent tag for cell membrane components, we estimated the cell boundary using the vRNP images. We assumed that fluorescent signal would be denser within the cell cytoplasm than in the surrounding area with cell-cell junctions subtly defined by ‘valleys’, curves of low signal density whose normal vectors point towards increasing density. The gradient of point density over the image was then used to segment individual cells from both the background and each other by the mean-shift algorithm [37]. Hand segmentation was also used to ensure accurate cell membrane segmentation. In most cases, the convex hull of all points within an identified cell region then defined the cell membrane. The nucleus was segmented through simple thresholding and smoothing.

### Point processes and patterns

Within each cell, the vRNP segments form a point pattern, **x** = {x_1_, x_2_, …, x_n_}, where *n* is the number of observed points and x_i_ is the 3-dimensional coordinate vector for point *i*. The point pattern is defined over a bounded region, *W*, the segmented cell cytoplasm. **x** is then viewed as a “realization” (an output) of some random point process **X**, the generating distribution for all patterns of the particular vRNP identity. The process **X** then represents the culmination of biological factors that determines vRNP segment location, eg. nuclear export, directed transport over the microtubule network, inhibition by other organelles, etc.

To model **X**, we first define its locational density, *f*, a function over the cytoplasm, where *f*(u) is the probability of observing a point at position **u**. The simplest model for this density is the Homogenous Poisson, characterized by complete randomness over space, or uniform probability:

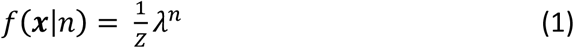

where *λ* determines point density, and *Z* a normalizing constant. In the point process literature, *λ* is referred to as *intensity*, but we refer to it as point density to avoid confusion with fluorescence intensity.

### Characterizing spatial randomness

Hypothesis testing for spatial randomness, comparing an observed pattern to an expectation under a Poisson assumption, was used as an initial motivation for further modeling. Ripley’s K-function describes the number of neighboring points within a given distance (*r*) of each observed point in a pattern, a measure of clustering or inhibition of points within space. Under a Poisson assumption,

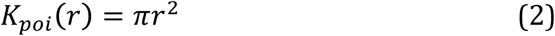

The expected K-function value can then be used to measure the difference between a given pattern and that of a Homogenous Poisson with an equal number of points. To assess this for observed point patterns, we used the test statistic

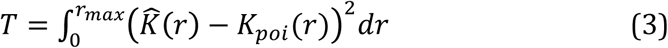

Where 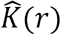 is the estimated K-function for a given pattern and *r* a radius defining the point neighborhood. To provide a background distribution for this statistic for a given cell geometry, we generated 100 samples from a Homogenous Poisson process defined over its cell cytoplasm and calculated the test statistic. We then obtained a p-value for the hypothesis that an observed distribution was drawn from a homogenous Poisson process:

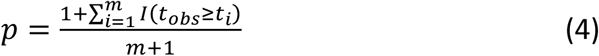

where *m* is the number of samples drawn. The p-values were averaged over all cells for a given vRNP.

### Modeling spatial dependence on cellular structures

An Inhomogenous Poisson Process, in its simplest form, is characterized by a spatially dependent locational density. Per point “factors” (e.g., distance to cell membrane) are used to determine the probability of a point occurring at a given location. As above, let **X** be some point process with realization **x**, and define *s*(**x**), an *n* × *k* matrix where each *s*_i,j_ is some factor *j* that can be calculated for each point x_i_. The Inhomogenous Poisson point density function is:

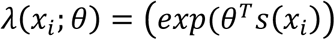

where θ is the k-dimensional parameter vector. The likelihood of the model given point data is then defined by:

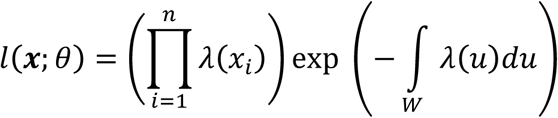

To quantify vRNP dependence on cellular structures, we defined the first factor (*f*_*1*_) as the distance to the nearest point on the nuclear membrane, the distance to the nearest point on the cell membrane, or the ratio of cell to nuclear distance.

### Modeling spatial dependence between vRNPs

The second factor, minimum inter-pattern point distance, was used to explore whether there is vRNP-vRNP interaction between vRNPs of different types. For each observed vRNP segment of type *l*, and some other vRNP point pattern of type *j*, the minimum inter-pattern point distance is:

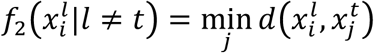

For each vRNP type, we use a ‘one depends on all’ schema. That is, each model with the same set of vRNPs is not equivalent but exclusively represents a single vRNP pattern (the primary pattern) depending on others (the secondary patterns). Models were constructed for every unique pair, triplet, and quadruplet of vRNP types.

For each model, all instances of its unique components were gathered, all images with tags for every vRNP in the model. Parameters of the model were then fit for each instance using the maximum pseudolikelihood [38]. The basic idea is that a given set of parameters can be used to calculate a value that is proportional to the likelihood (which is referred to as a pseudolikelihood) of observing a point at a given position, and the parameters can then be adjusted to give the highest total pseudolikelihood for available observations. Hold one out cross validation was used to assess the quality of fit over all instances yielding cross-validated pseudolikelihood estimated parameters and variances. These fitted parameters were then used to convert the pseudolikelihood to a true model likelihood by estimating the proportionality constant as previously described (19).

There is some variation in the number of images in which different vRNPs were visualized (Supplemental Table 1). This ranged from 4 for PB1 to 9 for M (the others were present 6 or 7 times). As pointed out by a reviewer, this difference raises the possibility that a vRNP that was imaged less frequently might be found to be underrepresented in the extent to which other segments were found to depend upon it. As seen in Table 3, this did not turn out to be the case, as PB1 and M showed similar average likelihoods and more vRNPs showed strong dependence on PB1 than on the others. Similarly, PB1 was not underrepresented in high scoring triples (Table 5).

### Interaction network assembly

We formulate the task of finding the most probable set of vRNA interactions yielding a full genome as a graph problem. First, let the set of nodes in the graph be every possible combination of vRNAs ranging in size from a single vRNA to all 8 segments. Directed edges in the graph connect subset to superset nodes. We weight each edge of the graph by the likelihood of the most likely model (chosen from models with only vRNP dependencies, and those with additional dependencies on either nuclear distance, cell membrane distance, or cell to nuclear distance ratio) that results in the superset and contains the subset as a primary or secondary segment (Figure 1). Edges that represent models that have not been observed remain unweighted.

From this graph, a set of interactions yielding the full genome is a tree with leaves as the vRNA singletons and root as the full 8-mer of vRNA segments. In these trees, we require each node to have at least two incoming edges, barring the leaf nodes. The set of incoming edges to a given node must emanate from nodes whose labels, when combined, exactly equal the given node. A set of incoming edges essentially ‘combines’ nodes, directly corresponding to the physical event of vRNP segments binding. The most likely set of interactions is also the most highly weighted tree. The problem of finding the most probable interaction network then reduces to finding the highest weighted tree in the graph. This problem can easily be solved through dynamic programming within a paradigm often used in evolutionary tree construction [24].

The likelihoods of models involving multiple vRNP segments can be taken as a measure of spatial dependence of primary segments on secondary segments. vRNP segments that truly interact are expected to be highly dependent either unidirectionally or bidirectionally. The cross validated model likelihoods can be used to weight groups of edges in the graph. Since the graph also has edges that were not observed in actual images (eg: any complex of more than 4 vRNAs), composite or inferred likelihoods were generated through recursion in a dynamic programming scheme. For each unobserved complex, we took the most likely set of observed models that could generate the complex.

For a given unobserved complex N = {N_1_,N_2_,N_3_,…},

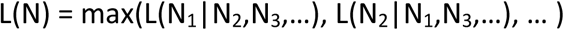

Then for some likelihood in the above, L(N_1_|N_2_) where N_1_ = (n^1^_1_, … n^1^_k_) and N_2_ = (n^2^_1_, … n^2^_l_),|N_1_| = k, |N_2_| = l,

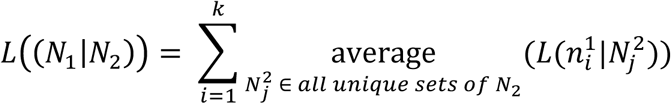

### Interaction network stability analysis

To assess the stability of the generated interaction network, two procedures of perturbation were performed: introduction of noise and removing certain high likelihood models. We simulated noise as a normal random distribution centered at 0 with varying standard deviations, termed the noise levels. Under increasing noise levels, each observed model likelihood was amended with a noise value drawn from this distribution prior to interaction network construction. For each noise level, we simulated 100 trials and tallied the proportion of output trees that contained each possible edge. Trees that align with the non-noisy assembly signal stability while highly variable trees signal instability.

### Reproducible research archive

All images, derived data and source code will be available upon publication at http://murphylab.cbd.cmu.edu/software.

---

**Supp. Table 1:** FISH imaging probe sets. The leftmost column is the experiment label with the right 4 columns showing which vRNA segments were tagged. Each experiment also used DAPI staining.

| Reaction | Probe 1 | Probe 2 | Probe 3 | Probe 4 |
| --- | --- | --- | --- | --- |
| A | M | HA | PB2 | NS |
| B | PB1 | PB2 | PA | NP |
| C | PB1 | M | HA | NS |
| D | M | PA | HA | NS |
| E | M | HA | NA | NP |
| F | PB1 | PB2 | NA | NS |
| G |  | PA | NA |  |
| H |  |  | NP | NS |
| I | PB1 | M | PB2 | PA |
| J |  | M | NA | NP |
| K |  | PB2 | HA | NP |
| L | M | HA | NA | NS |
| M | M | NA | HA | PA |
| N | M | PB2 | NP |  |
| Rxn 1 | M | PA | NA | NS |

**Supp. Table 2:**
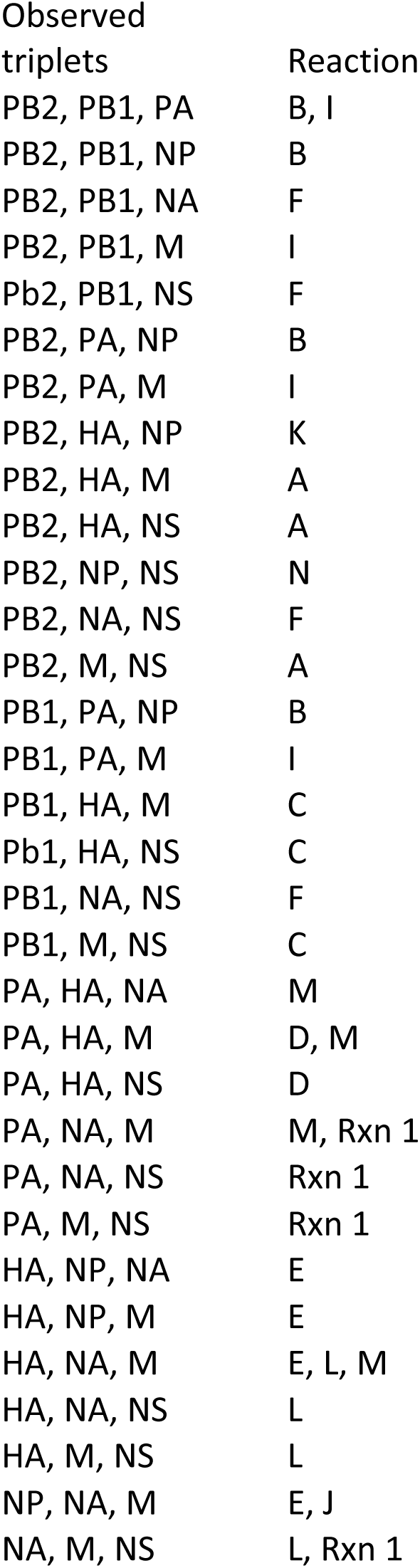
Observed triplets within the reaction set. 32 of 56 possible triplets were observed.

**Supp. Table 3:** Cross validated log-variances for pairwise interaction models shown in Table 3. (large)

**Supp. Table 4:** Variance in the cross validated negative log-likelihoods for pairwise interaction models with cell-to-nuclear distance ratio shown in Table 4. The row label indicates the independent variable, the vRNP upon which a model predicting the column label (dependent variable) was constructed.

|  | PB2 | PB1 | PA | HA | NP | NA | M | NS |
| --- | --- | --- | --- | --- | --- | --- | --- | --- |
| PB2 |  | 0.0552 | 0.0072 | 0.0310 | 0.0086 | 0.0028 | 0.0025 | 0.0919 |
| PB1 | 0.0172 |  | 0.0025 | 0.0012 | 0.0010 | 0.0002 | 0.0012 | 0.0067 |
| PA | 0.0091 | 0.0055 |  | 0.0054 | 0.0035 | 0.0061 | 0.0055 | 0.0103 |
| HA | 0.0175 | 0.0018 | 0.0011 |  | 0.0052 | 0.0064 | 0.0119 | 0.0146 |
| NP | 0.0489 | 0.0006 | 0.0015 | 0.0029 |  | 0.0060 | 0.0079 | 0.0003 |
| NA | 0.0001 | 0.0002 | 0.0023 | 0.0561 | 0.0143 |  | 0.0066 | 0.0025 |
| M | 0.0146 | 0.0022 | 0.0064 | 0.0146 | 0.0104 | 0.0037 |  | 0.0068 |
| NS | 0.0033 | 0.0068 | 0.0083 | 0.0038 | 0.0004 | 0.0011 | 0.0042 |  |

**Supp. Table 5:** Cross validated log-likelihoods for all observed triplets. (large)

**Supp. Table 6:** Cross validated log-likelihoods for all observed quadruplets. (large)

